# Arabidopsis WALL-ASSOCIATED KINASES are not required for oligogalacturonide-induced signaling and immunity

**DOI:** 10.1101/2024.04.15.589471

**Authors:** Laura Herold, Chenlei Hua, Bruce Kohorn, Thorsten Nürnberger, Thomas DeFalco, Cyril Zipfel

## Abstract

Carbohydrate-based cell wall signaling impacts plant growth, development, and stress responses; however, how cell wall signals are perceived and transduced remains poorly understood. Several cell wall breakdown products have been described as typical damage-associated molecular patterns (DAMPs) that activate plant immunity, including pectin-derived oligogalacturonides (OGs). Receptor kinases (RKs) of the WALL-ASSOCIATED KINASE (WAK) family have been shown to bind pectin and OGs, and were previously proposed as OG receptors. However, unambiguous genetic evidence for the role of WAKs in OG responses is lacking. Here, we investigated the role of Arabidopsis WAKs in OG perception using a novel deletion mutant of the clustered *WAK* family. Using a combination of immune assays for early and late pattern-triggered immunity (PTI), we show that WAKs are dispensable for OG-induced signaling and immunity, indicating that they are not *bona fide* OG receptors.

## Introduction

Plants are exposed to myriad potential pests and pathogens, against which they have evolved sophisticated defense mechanisms. The plant cell wall acts as the initial physical barrier against invasion, and alterations in this structure intricately interact with the plant immune system (Dora *et al*., 2022; Wolf, 2022).

The plant cell wall is composed of cellulose, hemicellulose, pectin, polyphenolic lignin and a series of structural and enzymatically active proteins (Wolf, 2022; Cosgrove, 2023). Cell wall polysaccharides serve as extracellular sources for the generation of damage-associated molecular patterns (DAMPs) that are thought to be released upon mechanical damage or pathogen infection (Pontiggia *et al*., 2020). Several such carbohydrate DAMPs have been previously described, including cellulose-derived cellobiose and cellotriose, mixed linked glucans, and pectin-derived oligogalacturonides (OGs) (Bacete *et al*., 2018; Oelmüller *et al*., 2023). OGs are generated from demethylesterified pectins and represent the best studied pectin-derived cell wall breakdown products. OGs with a degree of polymerization (DP) 10-15 (OG_10-15_) have been shown to elicit canonical PTI signaling and confer plant protection against a range of pathogens (Bishop *et al*., 1981; Hahn *et al*., 1981; Ridley *et al*., 2001; De Lorenzo *et al*., 2011). More recently, shorter OGs such as trimers (GalA_3_/OG_3_) and tetramers have also been shown to trigger immune responses and defense (Davidsson *et al*., 2017; Liu *et al*., 2023; Xiao *et al*., 2024).

In the model plant *Arabidopsis thaliana* (hereafter, Arabidopsis), demethylated pectin was shown to directly bind the extracellular domain (ECD) of several WALL-ASSOCIATED KINASES (WAKs) (Decreux and Messiaen, 2005; Decreux *et al*., 2006; Kohorn *et al*., 2009; Liu *et al*., 2023). WAKs belong to a large family of receptor kinases (RKs) comprising 5 WAKs and at least 21 WAK-likes (WAKLs) (Verica and He, 2002) that are characterized by epidermal-growth factor (EGF)-like domains and a galacturonan-binding domain in their ECD (He *et al*., 1996; Verica and He, 2002; Kohorn *et al*., 2012; Stephens *et al*., 2022). WAK1 was the first identified RK physically linking the plasma membrane (PM) to the cell wall, as isolation from fractions of proteolytically digested cell walls indicated a strong interaction of WAK1 with the cell wall and native pectin *in vivo* (He *et al*., 1996; Anderson *et al*., 2001; Wagner and Kohorn, 2001). Further experiments suggested WAKs and their association with cell wall pectin are involved in cell expansion (Kohorn *et al*., 2006), and, potentially the response to pathogens (He *et al*., 1998; Kohorn *et al*., 2012). Later, WAK1 was also shown to bind pectin and OG_9-15_ with a high affinity *in vitro* (Decreux and Messiaen, 2005; Decreux *et al*., 2006; Kohorn *et al*., 2009), with the WAK1-ECD preferentially interacting with de-esterified pectin through a binding site formed by cationic amino acids (Decreux and Messiaen, 2005; Decreux *et al*., 2006).

A chimeric approach using the ECD of WAK1 fused to the intracellular domain of the leucine-rich repeat RK ELONGATION FACTOR-TU RECEPTOR (EFR) served as evidence for a proposed role for WAK1 in OG perception (Brutus *et al*., 2010). Although OG treatment of WAK1-EFR chimera-expressing plants induced an EFR cytosolic domain-mediated defense response, critical genetic evidence that WAKs are *bona fide* OG receptors is still lacking. Direct genetics of WAKs was previously hindered by the genetic clustering of the *WAK* family in Arabidopsis, the assumption that *wak1* null mutants were lethal, and expected functional redundancy among the five members of the WAK family (He *et al*., 1999; Brutus *et al*., 2010; Kohorn and Kohorn, 2012). Recently, however, a CRISPR deletion mutant for most of the chromosomal cluster carrying the five Arabidopsis *WAK* genes, *wakΔ*, was generated. This mutant was shown to be less sensitive to the bacterial flagellin-derived epitope flg22, chitin and OGs in terms of reactive oxygen species (ROS) production (Kohorn *et al*., 2021); suggesting that WAKs may generally regulate immune receptor complexes, rather than function specifically as OG receptors (Wang *et al*., 2020; Zhang *et al*., 2020). WAKs were also recently shown to be genetically required for GalA_3_-induced expression of the salicylic acid (SA) marker gene *PATHOGENESIS-RELATED 1 (PR1)*(Liu *et al*., 2023).

In this work, we directly investigated the genetic involvement of WAKs in OG-induced signaling in Arabidopsis. We generated a novel deletion of the entire *WAK1-5* region (*wakΔ2*) and tested this mutant for OG-induced responses. Surprisingly, we found that *wakΔ2* retained full responsiveness to OGs, as measured by both early and late outputs of immune signaling. In addition, *wakΔ2* plants were not affected in OG-induced resistance against both bacterial and fungal pathogens. Furthermore, we tested the genetic involvement of WAKs in response to flg22 and could observe that flg22-induced responses are not affected in the *wakΔ2* mutant. Together, our data indicate that WAKs are not genetically required for OG perception and ensuing immune signaling in Arabidopsis.

## Results

### Generation of the *wakΔ2* mutant

The Arabidopsis genome has five *WAK* genes located in cluster on chromosome 1 (FIGURE ***1***A). Recently, a partial deletion mutant was published, *wakΔ,* which lacks most of the cluster; however, this mutant still potentially expresses a fusion protein of the N-terminal region of WAK4 and the C-terminal region of WAK2 (FIGURE ***1***A) (Kohorn *et al*., 2021). While the *wakΔ* mutant showed partially impaired flg22, chitin and OG responsiveness, it suffered from the presence of this potential WAK4-WAK2 fusion protein. Therefore, to explore if WAKs are genetically required for OG-induced responses, we generated a novel mutant using CRISPR/Cas9 that has a 23-kb deletion (*wakΔ2*), in which all *WAK* genes are absent (**Error! Reference source not found.**A-D). Lack of *WAK1-5* expression in *wakΔ2* seedlings was confirmed using RT-qPCR (FIGURE ***1***D). In agreement with the previously published *WAK* deletion mutants (Kohorn *et al*., 2021; Liu *et al*., 2023), *wakΔ2* displayed no obvious growth phenotype when grown on soil (FIGURE ***1***E).

**FIGURE 1.**
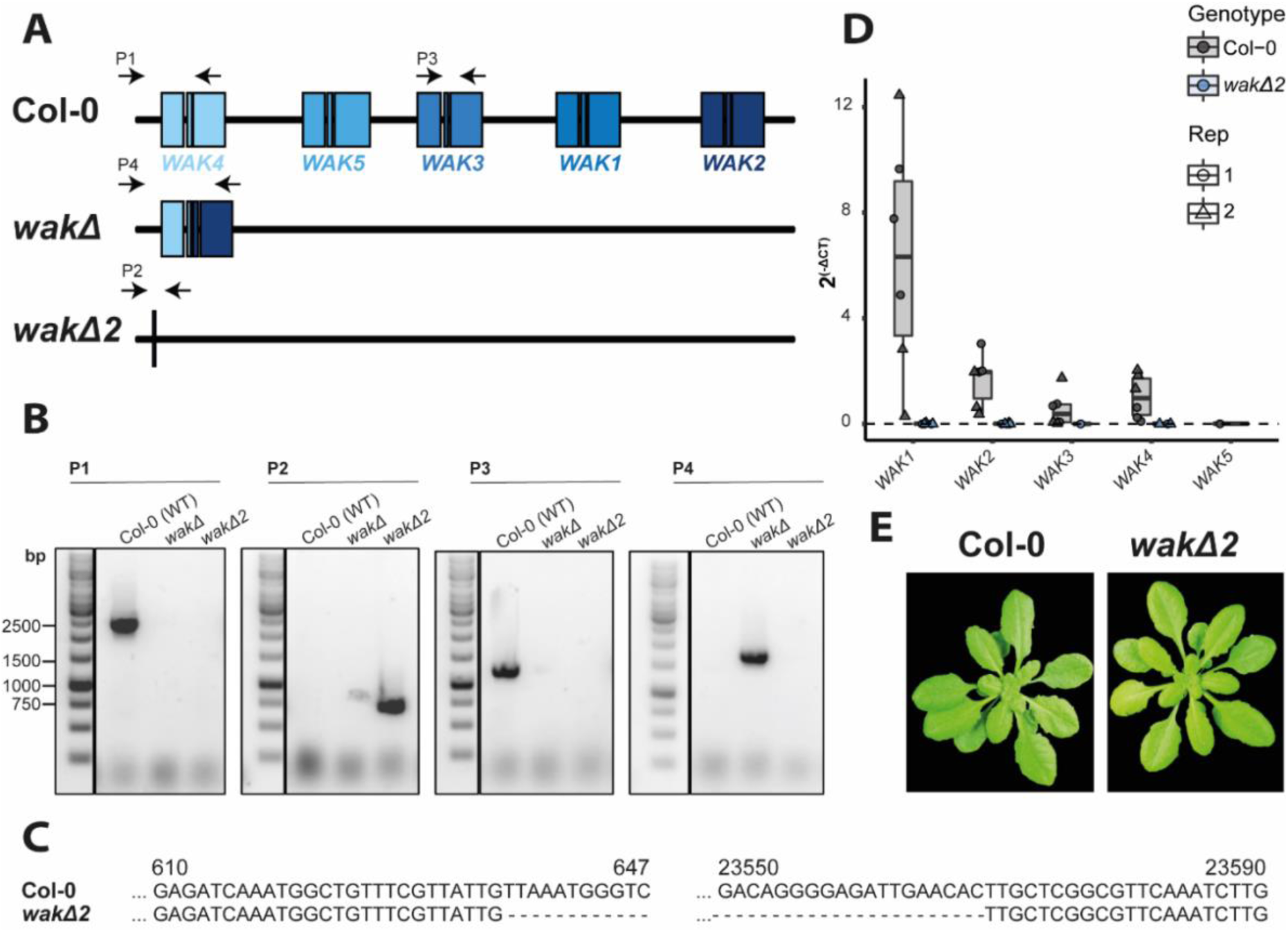
*wakΔ2* is a full *WAK* deletion mutant. **A)** Schematic representation of the genomic arrangement of *WAK1-5* in Arabidopsis. Middle cartoon shows the genomic deletion in *wakΔ* and the consequent fusion of WAK4 and *WAK2*. The lower cartoon shows the genomic region of the *WAK* cluster in *wakΔ2*. Black arrows indicate primer pairs (P) used in B. **B)** Genotyping gel of Col-0, *wakΔ* and *wakΔ2*. Ethidium bromide-stained PCR products for parts indicated in A on agarose gel. P1-P4 refer to the primer pairs shown in A. **C)** Sequencing results from the *wakΔ2* aligned against the 5’UTR of *WAK4* and part of the third exon of *WAK2* of Col-0. **D)** Transcript levels of *WAK1-5* in Col-0 and *wakΔ2* determined by RT-qPCR. RNA was extracted from 14-day-old Arabidopsis seedlings grown in liquid culture. Transcripts were normalized to the house-keeping gene *UBOX*. Three biological replicates per experiment (Rep) were used. **E)** Representative images of four-week-old Arabidopsis plants grown on soil. These experiments were performed two times.

### OG-induced immune signaling does not require the WAK family

Given that WAKs are proposed as receptors for OGs (Brutus *et al*., 2010), we investigated their genetic requirement for OG-induced immune responses using *wakΔ2.* Previous studies have extensively studied the immune responses in Arabidopsis triggered by exogenously applied OG_10-15_ including extracellular ROS production, mitogen-activated protein kinase (MAPK) activation, marker gene expression, ethylene production, callose deposition, seedling growth inhibition (SGI), and resistance against pathogens (Denoux *et al*., 2008; Davidsson *et al*., 2017; Gravino *et al*., 2017; Bjornson *et al*., 2021). Full loss-of-function mutants of a *bona fide* OG receptor should not be able to induce OG-induced responses, as shown for other ligand-perceiving receptors (Gómez-Gómez and Boller, 2000; Chinchilla *et al*., 2006; Zipfel *et al*., 2006; Miya *et al*., 2007; Yamaguchi *et al*., 2010; Cao *et al*., 2014; Ranf *et al*., 2015; Rhodes *et al*., 2021).

To investigate if early immune signaling induced by OGs is dependent on WAKs, we measured extracellular ROS production in leaves of 3- to 4-week-old Arabidopsis plants. Surprisingly and in contrast to previous results (Kohorn *et al*., 2021), OG_10-15_-induced ROS production was unaltered in *wakΔ2* in comparison to Col-0 grown in our conditions under short day (FIGURE 2A,B). In addition to ROS, OGs induce rapid and transient MAPK phosphorylation (Gravino *et al*., 2017). To determine if OG-induced MAPK activation is affected in the *wakΔ2*, MAPK phosphorylation was determined in Arabidopsis seedlings 5 and 15 min after elicitor treatment by western blot analysis using a commercial phosphorylation site-specific antibody. As with ROS production, OG_10-15_-triggered MAPK phosphorylation was unaltered in *wakΔ2* mutants (FIGURE 2C). In addition to OG_10-15_, OG_3_ (GalA_3_) was previously shown to trigger MAPK phosphorylation and WAKs have been shown to be required for OG_3_-induced *PR1* expression (Davidsson *et al*., 2017; Liu *et al*., 2023). We therefore additionally investigated if OG_3_-induced MAPK phosphorylation is dependent on WAKs. OG_3_-induced MAPK phosphorylation is comparatively weak but was still induced in *wakΔ2* mutant plants (FIGURE 2D). OGs were also previously shown to induce synthesis of ethylene in Arabidopsis seedlings (Ferrari *et al*., 2008; Brutus *et al*., 2010; Gravino *et al*., 2015). In line with other early induced PTI pathways, OG_10-15_-induced ethylene production was not compromised in *wakΔ2* mutants (FIGURE 2E). Together, these results indicate that WAK1-5 are not required for OG-induced early immune outputs and thus the signaling initiation of OG-induced responses.

**FIGURE 2.**
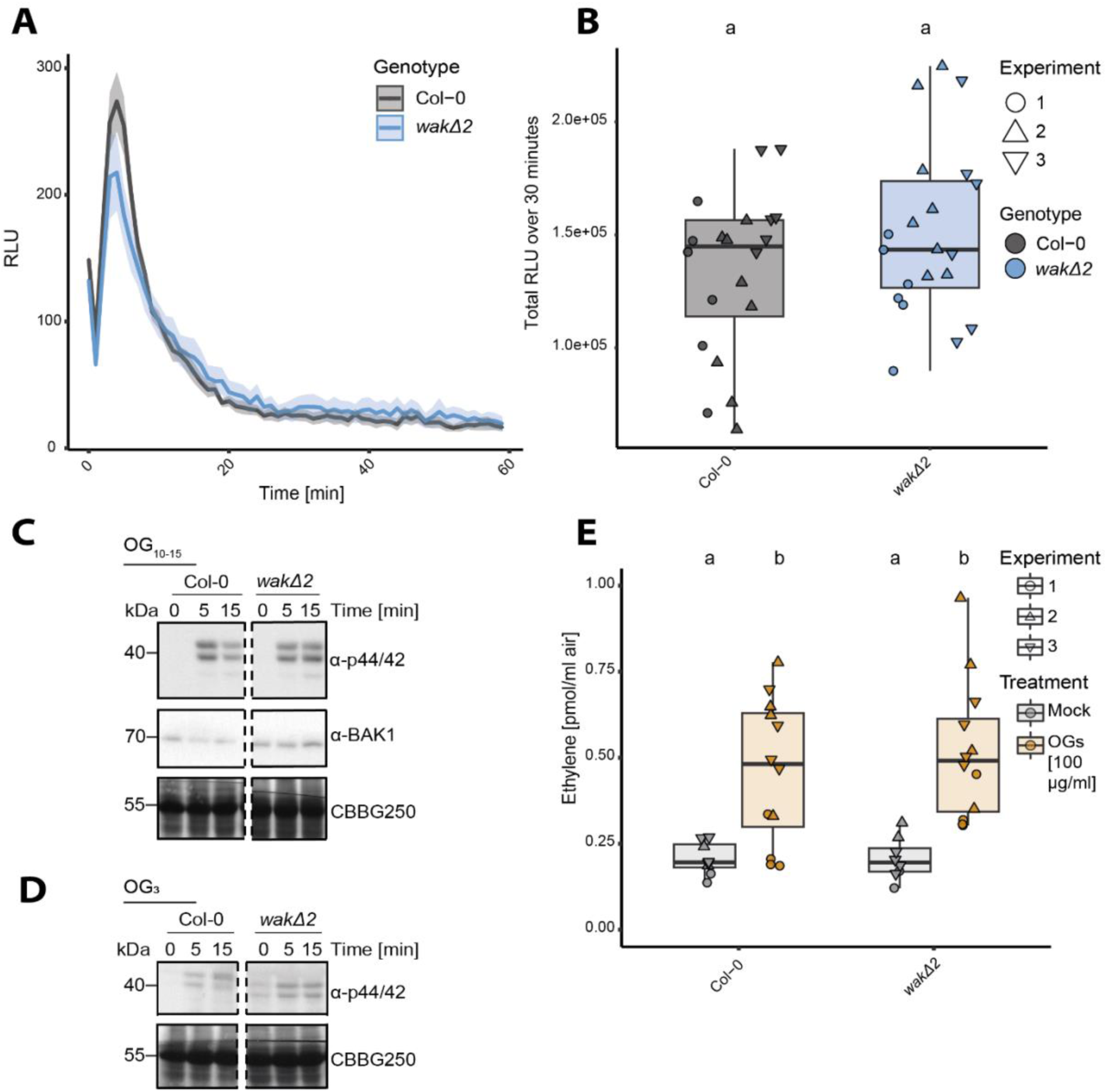
WAKs are not required for OG-induced early immune responses. **A-B)** ROS production in response to OG_10-15_ in leaf discs of 3- to 4-week-old Arabidopsis plants (n = 12 leaf discs of 6 plants). 100 µg/mL – 1 mg/mL of OG_10-15_ were used as concentration dependent of the experiment. Mean ± standard errors are plotted. RLU=relative luminescent units. **A)** Representative graph of the kinetics of one replicate. 1 mg/mL OG_10-15_ was used as concentration. **B)** Values are means of total photon counts over 30 min. Data from three independent experiments (Rep) are shown. Shapes indicate different replicates. Outliers are included in statistical analysis. Statistical test: Kruskal-Wallis test (p < 2.62*10^-08^), Dunn’s post-hoc test with Benjamin-Hochberg correction (p <= 0.05). Groups with like letter designations are not statistically different. **C-D)** MAPK activation assay with 2-week-old seedlings in response to 100 µg/mL OG_10-15_ (C) or 100 µg/mL OG_3_ (D). Samples were collected 0, 5 and 15 min after elicitation as indicated. Blot was probed with α-p44/42 and α-BAK1 was used as loading control. CBB=commassie brilliant blue was used as loading control as well. **E)** Ethylene accumulation after treatment with 100 µg/mL OG_10-15_ or water as control in Arabidopsis seedlings. Box plots represent means ± SE of three replicates. Equal letters at the top of the panel indicate p > 0.05, two-way ANOVA and a post hoc Tukey test. These experiments were performed three times.

Aside from rapid signaling, PTI additionally involves longer-term responses such as callose deposition (Beffa *et al*., 1996; Luna *et al*., 2011; Wang *et al*., 2021). To investigate the requirement of WAKs at later stages of OG-induced responses, OG_10-15_-induced callose deposition was measured in leaf discs of Col-0 and *wakΔ2* twenty-four hours after infiltration of either water or 100 µg/mL OG_10-15_. OG_10-15_-induced callose deposition in both Col-0 and *wakΔ2* (FIGURE 3 3A,B). As is true of many elicitors, both OG_3_ and long OGs can inhibit plant growth (Davidsson *et al*., 2017). Arabidopsis seedlings grown in the presence of OG_10-15_ showed a significant growth inhibition in comparison to mock-treated seedlings; however, no difference could be observed between Col-0 and *wakΔ2* (FIGURE 3 3C). Another long-term measurement of plant immune signaling is the production of SA and ensuing signaling, which can be inferred through the accumulation of the PR1 marker protein by immunoblotting (Tsuda *et al*., 2009; Zhang and Li, 2019; Bender *et al*., 2021). OG_10-15_ and flg22 induced robust PR1 accumulation twenty-four hours after elicitor infiltration into leaves of Col-0 plants. Both flg22 and OG_10-15_-induced PR1 accumulation was not affected in *wakΔ2* (FIGURE *3* 3D). OG_3_ induced very weak PR1 accumulation; however, no difference in PR1 accumulation could be detected between Col-0 and *wakΔ2* (FIGURE 3 *3*D). Collectively, these results indicate that WAKs are not required for OG-induced immune signaling.

**FIGURE 3.**
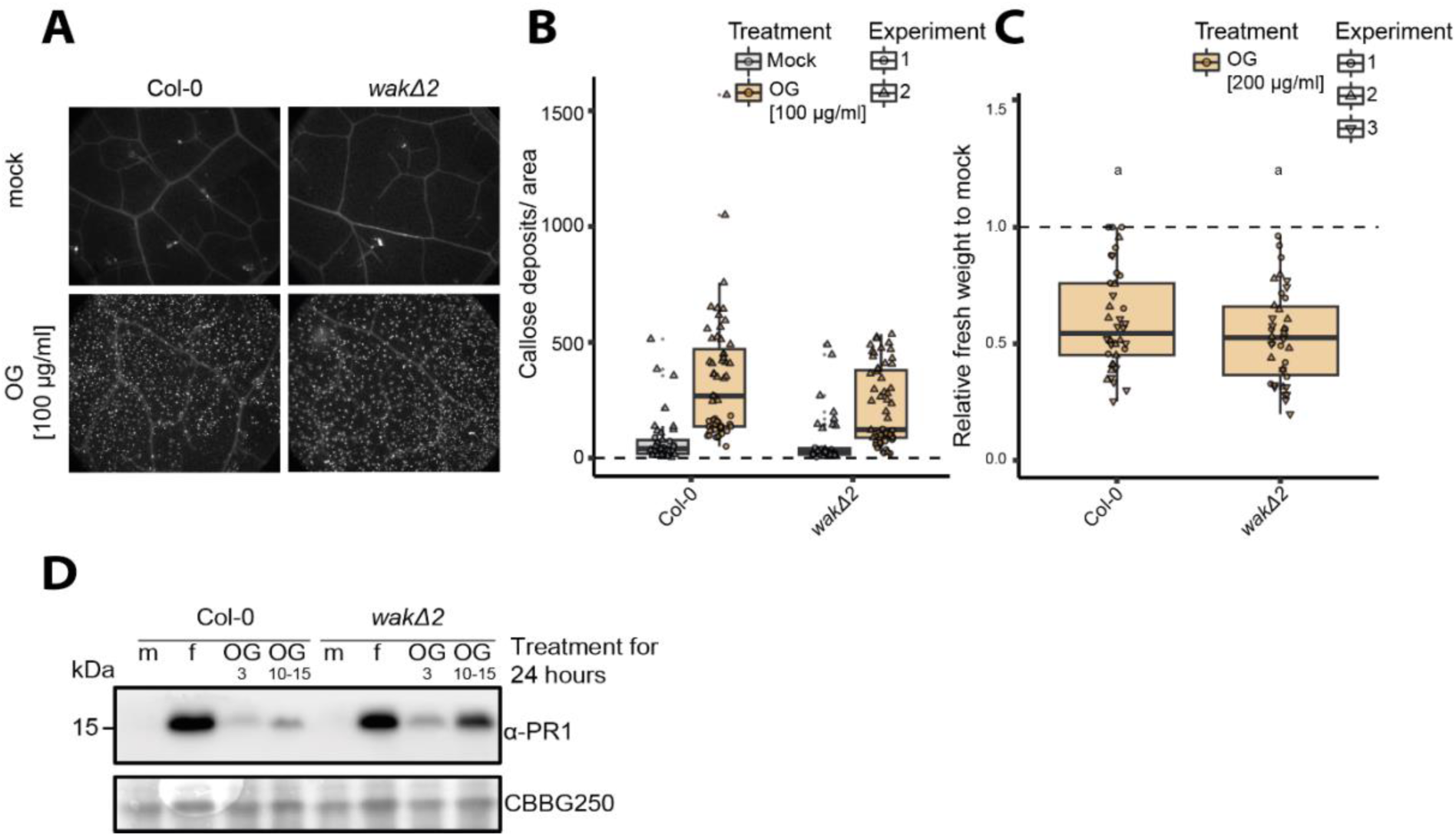
3 WAKs are not required for OG-induced late immune responses. **A-B)** Callose deposition visualized by aniline blue staining in response to 100 µg/mL OG_10-15_ or water 24 hours after infiltration into leaves of 3- to 4-week-old Arabidopsis plants. n=16-32 leaf discs from 4 different plants were taken per independent experiment. The experiment was performed two times with similar results. **A)** Representative images of OG-induced callose deposition in the presented genotypes stained with aniline blue. **B)** Callose deposits induced by OGs and water infiltration. **C)** Relative weight of seedlings grown in liquid media for 10 days in the presence of 200 µg/mL OG_10-15_ or in the absence of neither (mock). Means ± SE are shown with individual values for each plant and experiment (n = 12-14 seedlings per experiment). Outliers are included in statistical analysis. Equal letters at the top of the panel indicate p > 0.05, one-way ANOVA and a post hoc Tukey test. Groups with like letter designations are not statistically different. The experiment was repeated three times with similar results. **D)** PR1 accumulation assessed by immunoblotting with PR1 antibodies. Leaves from 3-week-old Arabidopsis plants were infiltrated with water (m = mock), 1 µM flg22 (f) or 100 µg/mL OG_10-15_ or 50 µg/mL OG_3_ and harvested after 24 hours. The experiment was repeated three times with similar results.

### WAKs are not required for OG-induced immunity

OGs have been shown to induce protection against the necrotrophic fungus *Botrytis cinerea,* the necrotrophic bacterium *Pectobacterium carotovorum* and the hemibiotrophic bacterium *Pseudomonas syringae* (Davidsson *et al*., 2017; Gravino *et al*., 2017; Howlader *et al*., 2020). To investigate if WAKs are required for OG-induced immunity, we drop-inoculated Arabidopsis Col-0 and *wakΔ2* leaves with *B. cinerea* conidia 24 h after infiltration with water or 100 µg/mL OG_10-15_. Disease lesions on leaves were measured 48 h post inoculation. Plants pre-treated with water showed significantly larger lesions sizes in both Col-0 and *wakΔ2* than plants that were pretreated with OG_10-15_ (FIGURE 4 *4*A,B). OG-induced protection against *B. cinerea* was not affected in *wakΔ2* plants in comparison to wild-type plants. OG induced protection against *P. syringae* was similarly unaltered in *wakΔ2* in comparison to Col-0 (FIGURE 4 *4*C). Overall, these results indicate that WAKs are not required for OG-induced immunity against these necrotrophic or hemi-biotrophic pathogens.

**FIGURE 4.**
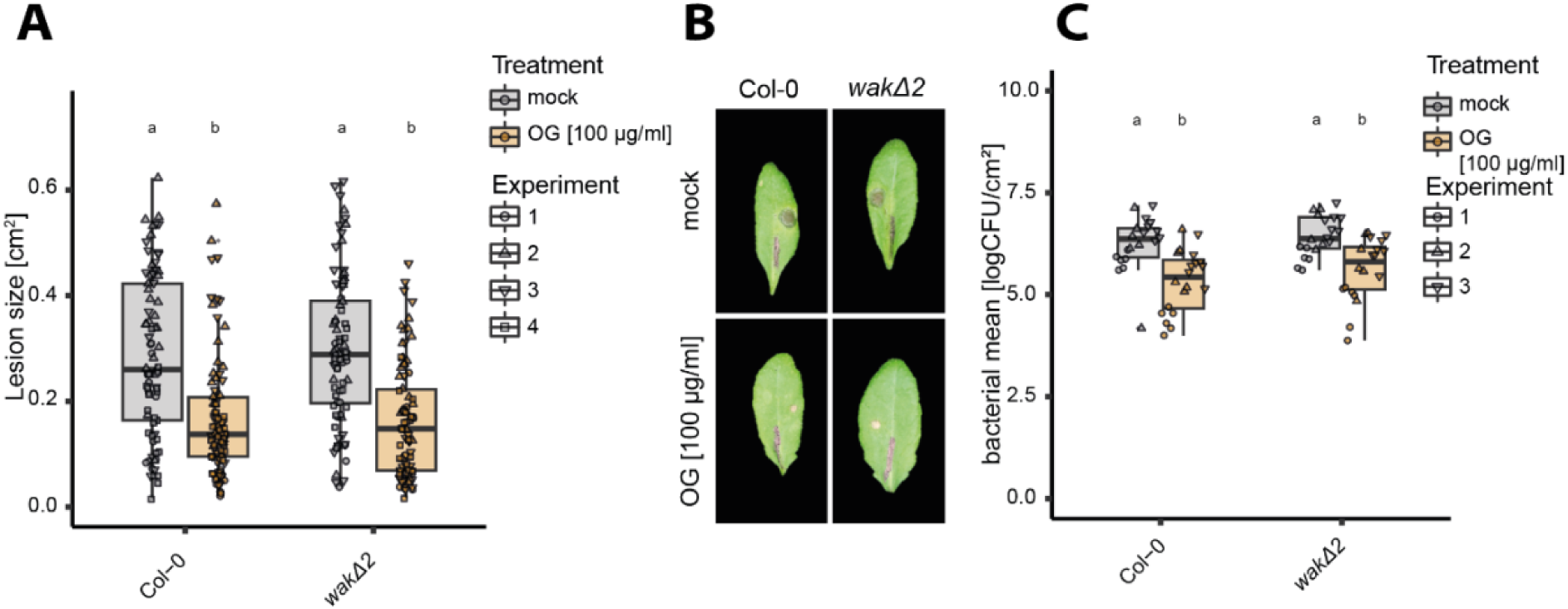
4 OG-induced immunity is not affected in *wakΔ2*. **A-B**) OG-induced resistance against *B. cinerea*. 4- to 5-week-old Col-0 or *wakΔ2* plants were infiltrated with water or 100 µg/mL OG_10-15_ 24hours prior drop-inoculation with *B. cinerea* strain BMM spores (5 µL; 5×10^5^ spores/mL). Lesion areas were measured 48 hours post inoculation. The experiment was performed four times. **A)** Quantification of lesion sizes. Results show mean ± SE (n = 18-24 per experiment). Equal letters at the top of the panel indicate p > 0.05, two-way ANOVA and a post hoc Tukey test. Groups with like letter designations are not statistically different. **B)** Representative images of OG-induced immunity in the different genotypes. Images were taken 48 hours post inoculation. **C)** OG-induced resistance against *P. syringae* pv tomato DC3000. Plants were pretreated with water or 100 µg/mL OG_10-15_ for 24 hours before infiltration with *P. syringae*. 48 hours after *P. syringae* infiltration, bacteria were extracted and plated. Results show means ± SE and individual data points from the three pooled experiments (n = 6 per experiment). Equal letters at the top of the panel indicate p > 0.05, two-way ANOVA and a post hoc Tukey test. Groups with like letter designations are not statistically different. The experiment was performed three times.

### WAKs do not play a significant role in immune signaling triggered by other elicitors

Aside from their role as potential OG receptors, WAKs were recently reported to function in immune signaling induced by bacterial flagellin in tomato and fungal chitin in cotton (Wang *et al*., 2020; Zhang *et al*., 2020). While in tomato only some flagellin-induced responses involved WAKs, e.g. callose deposition and anti-bacterial immunity, *Gh*WAK7A was broadly required for full responsiveness to fungal chitin but not to OGs in cotton (Wang *et al*., 2020; Zhang *et al*., 2020). In line with those observations, the Arabidopsis *wakΔ* mutant showed a reduction in ROS production induced by flg22, chitin and OGs (Kohorn *et al*., 2021). Intrigued by these findings, we also tested whether flg22-induced responses are affected by the full deletion of WAKs in Arabidopsis. In contrast to previous results, flg22-induced ROS production in leaves of 3- to 4-week-old Arabidopsis plants were not affected in *wakΔ2* in comparison to Col-0 under our conditions (FIGURE 5A,B). As expected, flg22-induced ROS production was dependent on the receptor FLAGELLIN-SENSING 2 (FLS2) and its co-receptor BRASSINOSTEROID-INSENSITIVE 1 (BRI1)-ASSOCIATED KINASE 1 (BAK1). In line with this, flg22-induced MAPK activation, ethylene production and induced resistance against *P. syringae* were not reduced in *wakΔ2* in comparison to Col-0 (FIGURE 5C-E). These results indicate that the deletion of WAKs does not affect flg22-induced responses under our growth conditions.

**FIGURE 5.**
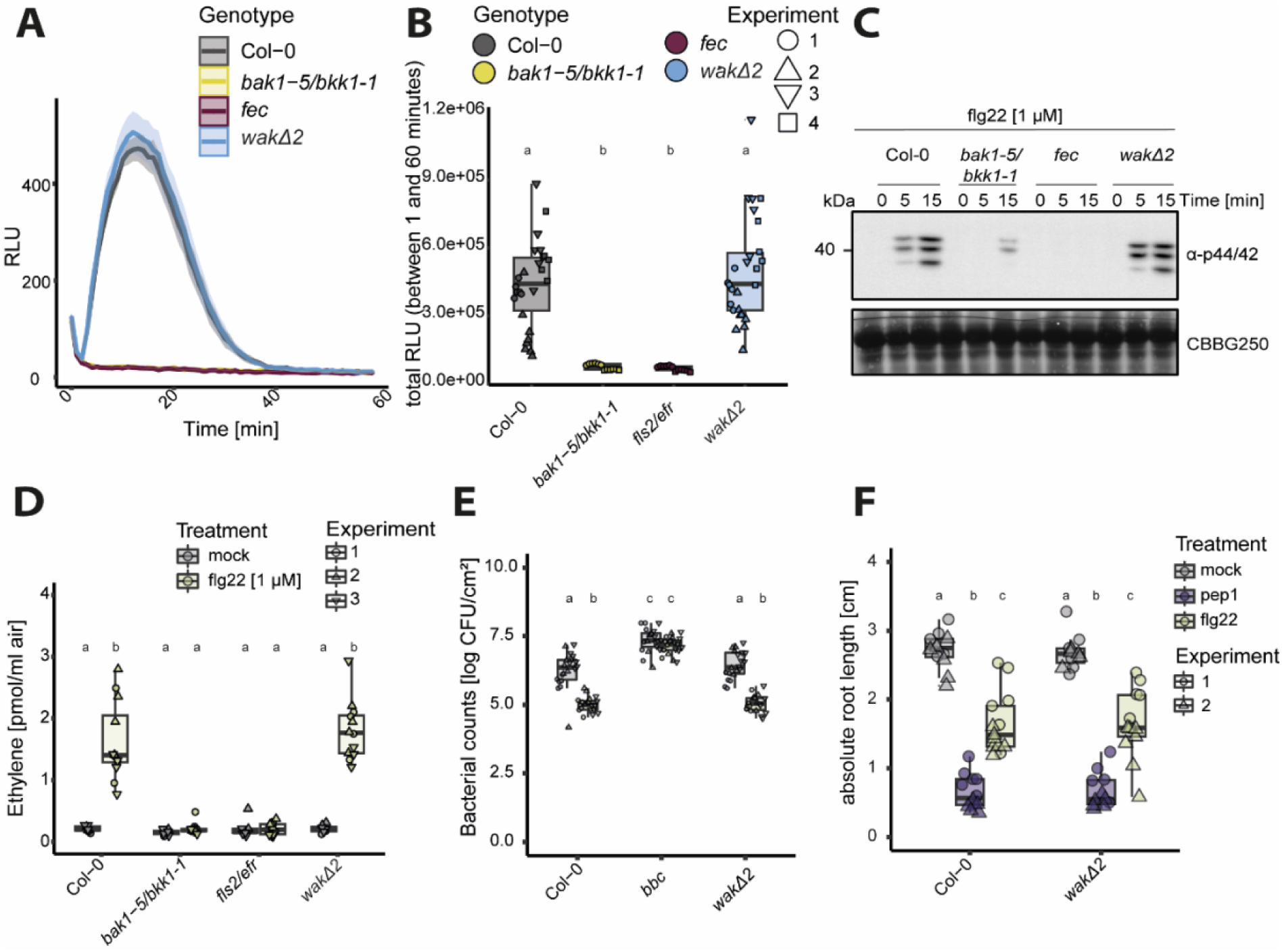
flg22-induced immunity is not affected by the loss of WAK1-5. **A-B)** ROS production in leaf discs of 3- to 4-week-old plants using 100 nM flg22 in Col-0, *bak1-5/bkk1-1*, *fls2/efr/cerk1* (*fec*) and *wakΔ2*. The experiment was repeated at least three times (A-B). Mean ± standard errors are plotted. RLU=relative luminescent units. A) Kinetics of three representative independent replicates over 40-60 minutes. B) Values are means of total photon counts over 60 minutes as stated in the graph. Individual data points show ROS production in individual plants (n = 6-8 plants with each two leaf discs). Outliers are included in statistical analysis. Kruskal-Wallis Test flg22 (p-value = 9.364.10^-15^), Dunn’s post-hoc test with Benjamin-Hochberg correction (p <= 0.05). Groups with like letter designations are not statistically different. **C)** MAPK activation assay with 2-week-old seedlings in response to 1 µM flg22. Samples were collected 0, 5 and 15 min after elicitation as indicated. Blot was probed with α-p44/42. CBB = commassie brilliant blue was used as loading control as well. **D)** Ethylene accumulation after treatment with 1 µM flg22 or water as control in Arabidopsis seedlings. Box plots represent means ± SE of three replicates. Equal letters at the top of the panel indicate p > 0.05, two-way ANOVA and a post hoc Tukey. **E)** OG-induced resistance against *P. syringae* pv. tomato DC3000. Plants were pretreated with water or 1 µM flg22 for 24 hours before infiltration with *P. syringae*. 48 hours after *P. syringae* infiltration, bacteria were extracted and plated. Results show means ± SE and individual data points from the three pooled experiments (n = 6 per experiment). Equal letters at the top of the panel indicate p > 0.05, two-way ANOVA and a post hoc Tukey test. Groups with like letter designations are not statistically different. **F)** Primary root length of Col-0 and *wakΔ2* seedlings. Plants were grown on ½ MS +1 % sucrose plates for 5 days and then transferred to liquid ½ MS +1 % sucrose without elicitor (mock), 10 nM pep1 or 100 nM flg22. Root-growth was determined after 4 days in liquid culture. 6 plants were measured per experiment. Values correspond to length of each root in cm. Equal letters at the top of the panel indicate p > 0.05, two-way ANOVA and a post hoc Tukey test. All experiments were performed three times with similar results, only primary root length was only measured twice.

The *wakΔ2* mutant had no obvious growth phenotype when grown on soil (FIGURE 1E). The only *wak*-related growth phenotype previously observed was reduced root length when seedlings were grown on MS medium lacking sucrose, most pronounced on 1/6 MS (Kohorn *et al*., 2006, 2021). Therefore, to investigate if elicitor-induced root-growth inhibition is affected in the *wakΔ2* mutant, both Col-0 and *wakΔ2* plants were grown in the presence of 10 nM *At*pep1, 100 nM flg22 or without elicitor for 5 days. Elicitor-induced root-growth inhibition was similar in Col-0 and *wakΔ2* in the same extent to Col-0 for both flg22 and *At*pep1 (FIGURE 5F).

## Discussion

PTI is achieved by the recognition of diverse elicitor molecules as ligands for plasma membrane-resident pattern recognition receptors (PRRs) (DeFalco & Zipfel, 2021). Cell walls are the first layer of defense against invading pathogens, many of which have evolved arsenals of enzymatic and mechanical means to degrade or penetrate the cell wall (Bacete *et al*., 2018; Dora *et al*., 2022). Thus, the integrity of the cell wall needs to be carefully monitored by sensor proteins. Several RKs have been proposed as PRRs that perceive cell wall breakdown products, including WAKs based on their ability to interact with pectin and its breakdown products (He *et al*., 1996; Decreux and Messiaen, 2005; Decreux *et al*., 2006; Kohorn *et al*., 2009; Brutus *et al*., 2010). Yet, genetic evidence that WAKs function as *bona fide* OG receptor(s) was missing. Here, we have used the *wakΔ2* mutant, which lacks all five members of the WAK family, to demonstrate that none of the WAKs are required for responses to either short or long chain demethylated OGs in Arabidopsis.

Previously, the galacturonan-binding domain of WAKs was shown to bind both pectins and demethylesterified OG_9-15_ (Decreux and Messiaen, 2005; Decreux *et al*., 2006). Additionally, chimeric WAK-EFR receptors were able to induce EFR-like responses upon OG-treatment (Brutus *et al*., 2010). While our results indicate that WAKs are not genetically required for OG-induced responses, they do not contradict the ability of WAK ECDs to bind pectins or pectin breakdown products. Interestingly, the ECD of the malectin-like RK FERONIA was also recently reported to bind to pectin and pectin breakdown products (Feng et al. 2018; Tang et al. 2022; Lin et al. 2022), suggesting that this biochemical property might be true for several cell wall-anchored RKs without necessarily functioning as the true receptors for these carbohydrates.

OG_10-15_ were suggested to be produced during pathogen infection and to subsequently induce immune signaling (Ferrari *et al*., 2013; Xiao *et al*., 2024). Although demethylesterified OG_10-15_ are active as elicitors, recent evidence challenges their production *in planta* as most Ogs produced during infection with *B. cinerea* or *Fusarium oxysporum* were acetyl- and methylesterified (Voxeur *et al*., 2019; Gámez-Arjona *et al*., 2022). While pectic fractions of various sizes and modifications show elicitor activity, the complexity of those *in planta*-produced Ogs as well as the profile of crude extracts produced in the lab complicates the attribution of individual OG species to the elicitor activity (Liu *et al*., 2023). Regardless of the exact nature of *in planta* OG species, WAKs have been previously proposed as the receptors for demethylesterified OG_10-15_ based on *in vitro* binding studies and chimeric approaches, and we are here unable to confirm any corresponding genetic requirement for WAKs in OG_10-15_-induced signaling. Interestingly, electrostatic analysis of the WAK1 ECD predicted by Alphafold revealed a negatively charged galacturonan-binding domain at apoplastic pH, contradicting the suggested binding of polyanionic de-esterified pectins (Lee and Santiago, 2023).

While our data demonstrate that members of the WAK family are not required for OG-induced responses, it is possible that quantitative phenotype(s) are masked by persistent functional redundancy. WAKs are characterized by a galacturonan-binding domain and many WAKs contain one or more copies of EGF-like domains in their ECD (Verica and He, 2002; Stephens *et al*., 2022). In addition to 5 WAKs, there are at least 21 WAK-likes (WAKLs) in Arabidopsis. To date, clear evidence is missing that WAKLs are also able to bind cell wall fragments (Kohorn, 2016), with the exception of WAKL22/RESISTANCE TO FUSARIUM OXYSPORUM 1 (RFO1) and WAKL14 (Huerta *et al*., 2023; Ma *et al*., 2024). However, based on their phylogenetic relationship, WAKLs are obvious candidates to test for further genetic redundancy.

Aside from a role in OG perception, WAKs were recently suggested to be involved in the regulation of other RK complexes during immunity, indicating that they might serve as accessory RKs of PRR complexes. In tomato and cotton, WAKs interact with and positively regulate PRR complexes and are required for full responsiveness to the corresponding elicitors (Wang *et al*., 2020; Zhang *et al*., 2020). In Arabidopsis, the *wakΔ* mutant was less sensitive to multiple elicitors in terms of ROS production (Kohorn *et al*., 2021). In contrast with these previous results, no quantitative reduction in OG- or flg22-induced ROS production could be observed in the *wakΔ2* mutant. Although this difference is striking, it might further underline the role of WAKs as accessory RKs under certain growth conditions rather than OG-perceiving receptors. While WAKs appear to interact with multiple elicitor-perceiving RKs, the exact mechanisms by which WAKs regulate immunity seem to differ between plant species or different WAKs.

WAKs are found across land plants, with the WAK/WAKL family expanded in monocots (de Oliveira *et al*., 2014; Kanyuka and Rudd, 2019; Stephens *et al*., 2022; Zhang *et al*., 2023a; Ngou *et al*., 2024). Several *WAKs* or *WAKLs* have been identified as resistance genes and are required for basal resistance to pathogens in a variety of different crop plants (Diener and Ausubel, 2005; Zuo *et al*., 2015; Hurni *et al*., 2015; Hu *et al*., 2017; Saintenac *et al*., 2018; Bot *et al*., 2019; Larkan *et al*., 2020; Li *et al*., 2020; Stephens *et al*., 2022; Zhang *et al*., 2023b; Dai *et al*., 2024; Zhong *et al*., 2024). While diverse roles and mechanisms for WAKs in plant immunity have been proposed, a clear possibility is that WAKs perceive pathogen-derived molecules. Indeed, Arabidopsis WAK3 was recently shown to be required for immune responses induced by bacterial harpins (Held *et al*., 2024) indicating that WAKs might indeed perceive microbial molecules. Additionally, three WAKs have also been demonstrated to exhibit a gene-for-gene interaction with specific pathogenic effectors in crops (Stephens *et al*., 2022). The WAK proteins Stb6 and Rlm9 provide resistance against *Zymoseptoria tritici* isolates expressing AvrSbt6 in wheat and *Leptosphaeria maculans* expressing AvrLm5-9 in oilseed rape, respectively (Brading *et al*., 2002; Larkan *et al*., 2016; Larkan *et al*., 2020). While no direct interaction could be detected between these fungal effectors and corresponding WAK resistance proteins (Saintenac *et al*., 2018; Larkan *et al*., 2020), intriguingly, a direct interaction has been observed between the maize WAK protein Snn1 and the *Phaeosphaeria nodorum* effector protein *Sn*Tox1. Unlike most other WAKs studied thus far, Snn1 serves as susceptibility factor for *P. nodorum* leading to disease in Snn1-expressing plants (Liu *et al*., 2012; Stephens *et al*., 2022; Shi *et al*., 2023). Maize qRgls1**/**WAKL^Y^ was also recently shown to confer quantitative disease resistance against gray leaf spot caused by the fungi *Cercospora zeae-maydis* and *C. zeina* (Zhong *et al*., 2024). Notably, an aqueous extract of *C. zeina* hyphae and spores was sufficient to induce WAKL^Y^-dependent ROS production suggesting that WAKL^Y^ perceives a fungal ligand.

Altogether, there is emerging evidence that WAKs may perceive diverse molecules of microbial origin and orchestrate both broad-spectrum and race-specific resistance (Kanyuka and Rudd, 2019), which is consistent with our evidence that they do not function as *bona fide* OG receptor(s) in Arabidopsis. However, the mechanisms by which WAKs contribute to immunity and their true ligand(s) remain to be definitively characterized.

## Material and methods

### Plant growth

Arabidopsis seeds were surface-sterilized using ethanol, plated on 0.5 MS medium (1 % sucrose, pH 5.8, 0.9 % phytoagar), stratified for 48 hours in the dark at 4 °C, and grown at 22 °C under a 16-hour photoperiod (120 μmol * s^-1^*m^-2^ illumination). For assays in adult plants, including ROS production, pathogen infection and callose deposition, seedlings were transferred to soil after 7-10 days growth on plates. Plants were grown in short-day cycles (10 h light/14 h dark, 60 % humidity, 20 °C) for an additional 2-3 weeks. For assays with seedlings, including MAPK activation, RNA extraction, seedling growth inhibition and root growth inhibition, these were transferred 5 days after exposure to light to liquid MS and grown there for 10 days. Mutants were generated in the *A. thaliana* Columbia (Col-0) ecotype and primers for genotyping are found in SUPPLEMENTAL TABLE 1.

### CRISPR-Cas9 mutagenesis

The *WAK4* and *WAK2* oligonucleotides used as templates for SgRNA-targeted sites (GCTGT TTCGTTATTGTTAAATGG) 432 bp 5’ to the *WAK4* ATG start codon, and (GGGGAGATTGAACAC TTGCTCGG) 77 bp 5’ to the *WAK2* stop codon were each cloned into pSkAtu26 (Feng et al., 2013). These two expression Sg cassettes were then cloned into pCambia1302 that also had a pOLE1-RFP cassette inserted into the ASN718 site by PCR cloning (Shimada et al., 2010).

The T1, RFP^+^ (expressing linked CAS9 and sgRNAs) were screened for a deletion by PCR using primers flanking the deletion, and then T2 RFP^-^ plants were screened again by PCR to isolate a plant with a deletion but not expressing CAS9 or the sgRNAs. These isolates were self-crossed to generate a homozygous deletion.

### RNA extraction and real-time (RT) quantitative PCR analysis

Total RNA was extracted from 2-week-old liquid-grown seedlings. Total RNA was extracted using TRI reagent (Sigma-Aldrich). To remove genomic DNA, samples were treated with TURBO DNA-free Kit (Thermo Fisher Scientific). cDNA synthesis was performed using 1 µg of DNA-free RNA sample with RevertAid First Strand cDNA Synthesis Kit (Thermo Fisher Scientific) according to the manufacturer’s protocol. RT-qPCR analysis was performed using diluted cDNA as template for PowerUp SYBR Green (Applied biosystems) with the primers provided in SUPPLEMENTAL TABLE 1.

### MAPK activation

MAPK activation was performed as previously described (Mühlenbeck *et al*., 2023). Five-day-old seedlings were transferred into 24-well plates containing 1 mL of liquid 0.5 MS (1 % sucrose). Two seedlings per well were grown there for another 10-12 days. Seedlings were treated with 100 µg/mL OG_10-15_ (Elicityl, GAT114), 100 µg/mL OG_3_ (GalA3, Megazyme) or 1 µM flg22, and harvested at each time point as indicated in figure captions. Total proteins were extracted using extraction buffer (50 mM Tris pH 7.5 (HCl), 150 mM NaCl, 10 % (v/v) glycerol, 2 mM EDTA, 1 mM homemade PPI (equivalent to Sigma-Aldrich Protease-inhibitor cocktail P9599), 1 mM NaF, 1 mM sodium-orthovanadate, 2 mM sodium-molybdate, 4 mM sodium-tartrate, 1 % (v/v) IGEPAL CA630, 5 mM DTT). Proteins were analyzed by SDS-PAGE and immunoblotting using p44/42 MAPK antibody (Cell Signaling Technology).

### Seedling growth inhibition

Seedling growth inhibition assays was performed as previously described (Abarca *et al*., 2021). Briefly, 5-day-old Arabidopsis seedlings were transferred to 48-well plate with one seedling per well. Each well contained either 500 µL 0.5 liquid MS with or without 200 µg/mL OG_10-15_ (Elicityl, GAT114). After 10 days of growth in the presence of the respective elicitor, individual seedling weight as assessed using an analytical balance.

### Root growth Inhibition

Five-day-old seedlings were transferred from solid MS plates to 12-well plate with 6 seedling per well. Each well contained 4 mL of liquid MS supplemented with mock (sterile ddH_2_O), 10 nM *At*pep1 or 100 nM flg22. After 5 days of treatment, seedlings were transferred to MS plates and imaged. Root lengths were quantified with ImageJ.

### Ethylene production

Four- to six-week-old Arabidopsis leaves were cut into 3-mm slices and floated on water overnight. For each sample, three leaf slices were transferred to a 6-mL glass tube containing 200 µL MES buffer (pH 5.7), followed by adding either water control or the elicitor to a final concentration of 1 µM. Vials were closed with a rubber septum and ethylene production in the free air space was measured by gas chromatography (Shimadzu, GC-14A) after 3 hours of incubation.

### ROS production

Leaf-discs of 3- to 4-week-old plants were taken (4 mm Ø) and placed with the abaxial side down into a well of a white polystyrene 96-well plate containing 100 μL ddH_2_O and recovered overnight. The next day, the water was replaced by a solution containing 20 µg/mL horseradish peroxidase (HRP, sigma), luminol (17 µg/mL) and elicitor (100 nM for flg22, 100 µg/mL OG_10-15_ (Elicityl, GAT114), as stated). Luminescence was immediately measured for 60 minutes using a charge-coupled device camera (Photek Ltd, East Sussex UK).

### Callose deposition

Callose deposition assays were performed as described previously (Mason *et al*., 2020). Briefly, four leaves of 4- to 5-week-old plants were syringe-infiltrated with either mock (ddH_2_O), 1 µM flg22 or 100 µg/mL OG_10-15_ (Elicityl, GAT114). Twenty-four hours after infiltration, leaf discs were taken and collected in 24-well plates filled with 1 mL 100 % EtOH until completely destained. Leaf discs were equilibrated in 1 mL 67 mM K_2_HPO_4_ (pH 12) for 60 min. Afterwards, the tissue was stained using aniline blue (Acros Oganics) staining solution (0.01 % (w/v) aniline blue in 67 mM K_2_HPO_4_ (pH 12) for 60 min and washed in 67 mM K_2_HPO_4_ (pH=12) for 60 minutes. Stained tissue was mounted in mounting solution (80 % glycerol, 67 mM K_2_HPO_4_, pH 12) on microscope slides. Callose deposits were imaged using a Leica DM6000B and quantified in ImageJ.

### PR1 protein abundance

PR1 accumulation was assayed as previously described (Bender *et al*., 2021). Briefly, three leaves of 3-week-old plants were infiltrated with mock (sterile ddH_2_O), 1 µM flg22, 100 µg/mL OG_10-15_ (Elicityl, GAT114) or 100 µg/mL OG_3_ (GalA3, Megazyme). Twenty-four hours after infiltration, leaves were harvested in 1.5-mL tubes and snap-frozen in liquid nitrogen and pulverized. Extraction buffer (50 mM Tris pH7.5 (HCl), 150 mM NaCl, 10 % (v/v) glycerol, 2 mM EDTA, 1x plant protease inhibitor cocktail) was added and protein concentration was adjusted by Bradford assay. Normalized protein extracts were analyzed by SDS-PAGE (15 %) and immunoblotting using PR1-antibodies (Agrisera).

### Induced resistance against *Pseudomonas syringae*

Two leaves of 4- to 5-week-old plants were infiltrated with 1 µM flg22 or 100 µg/mL OG_10-15_ (Elicityl, GAT114) or mock (sterile ddH_2_O). Freshly restreaked *P. syringae* pv tomato DC3000 was grown in liquid Kings B overnight and refreshed in a subculture the next morning for additional 1-2 hours. Bacteria were infiltrated into pretreated leaves with an OD_600_ of 0.0002. Plants were covered for two days, after which 1 leaf disc was harvested per treated leaf (8 mm Ø) and pooled per plant. Leaf discs were ground in 10 mM MgCl_2_, thoroughly mixed and diluted in a 1:10 series until 1:10^-6^. Samples were plated on LB plates. After two days of growth at 28 °C, colony forming units were counted. Statistics were performed on log_10_ (CFU/cm^2^).

### Induced resistance against *Botrytis cinerea*

Four leaves of 4- to 5-week-old plants were infiltrated with 1 µM flg22 or 100 µg/mL OG_10-15_ (Elicityl, GAT114) or mock (sterile ddH_2_O) in the morning. The next day, spores of *B. cinerea* BMM were collected in sterile ddH_2_O and the spores were counted using a counting chamber. At least 1 hour prior infection, infection solutions were prepared with a final concentration 5×10^5^ spores/mL in 0.5 Potato Dextrose Broth and incubated at RT. Five microliters of the *Botrytis* infection solution were dropped on the adaxial site next to the middle vein. Plant solid trays were filled with water, covered with a lid, and sealed with parafilm to produce high humidity. After 2 days after infection at dimmed light, leaves were detached, images were taken, and lesion size was measured using Image J.

## Author Contributions

L.H. and C.H. performed the experiments and analyzed the data. B.K. generated the genetic material. T.N., T.A.D and C.Z designed and supervised the project. L.H. wrote the first draft of the manuscript. All authors contributed to the final version of the manuscript.

## Acknowledgments

We thank all the members of the Zipfel group for fruitful discussions during the project. We thank also Jiashu Chu, John Haidoulis and Jana Ordon for feedback on the manuscript. L.H. was funded by a Zurich-Basel Plant Science Center-Syngenta Fellowship. B.D.K. was supported by National Science Foundation grant IOS 1556057. T.N. was supported by DFG-TRR356 (B5). T.A.D. was supported by a Discovery Grant from the Natural Sciences and Engineering Council of Canada (NSERC RGPIN-2023-04222). C.Z. was supported by the University of Zürich and the Swiss National Science Foundation grants no. 31003A_182625 and 310030_212382.

## Supplemental table

**SUPPLEMENTAL TABLE 1.**
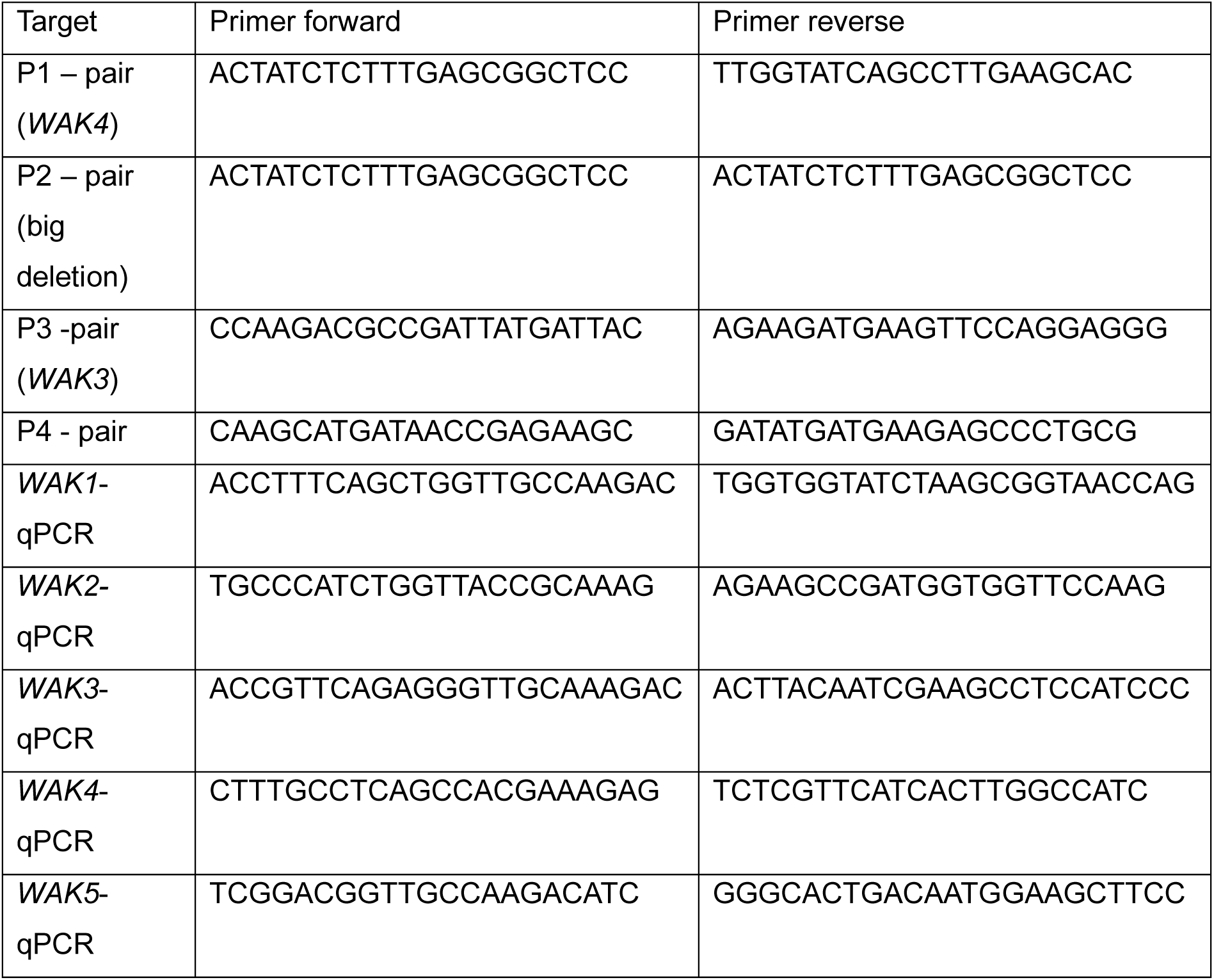
Primers used in this study.

## References

Abarca A, Franck CM, Zipfel C. 2021. Family-wide evaluation of RAPID ALKALINIZATION FACTOR peptides. Plant Physiology 187, 996–1010.

Anderson CM, Wagner TA, Perret M, He ZH, He D, Kohorn BD. 2001. WAKs: cell wall-associated kinases linking the cytoplasm to the extracellular matrix. Plant Molecular Biology 47, 197–206.

Bacete L, Mélida H, Miedes E, Molina A. 2018. Plant cell wall-mediated immunity: cell wall changes trigger disease resistance responses. The Plant Journal 93, 614–636.

Beffa RS, Hofer RM, Thomas M, Meins FJ. 1996. Decreased Susceptibility to Viral Disease of [beta]-1,3-Glucanase-Deficient Plants Generated by Antisense Transformation. The Plant Cell 8, 1001–1011.

Bender KW, Couto D, Kadota Y, et al. 2021. Activation loop phosphorylation of a non-RD receptor kinase initiates plant innate immune signaling. Proceedings of the National Academy of Sciences of the United States of America 118, (38):e2108242118.

Bishop PD, Makus DJ, Pearce G, Ryan CA. 1981. Proteinase inhibitor-inducing factor activity in tomato leaves resides in oligosaccharides enzymically released from cell walls. Proceedings of the National Academy of Sciences of the United States of America 78, 3536–3540.

Bjornson M, Pimprikar P, Nürnberger T, Zipfel C. 2021. The transcriptional landscape of Arabidopsis thaliana pattern-triggered immunity. Nature Plants 7, 579–586.

Bot P, Mun B-G, Imran QM, Hussain A, Lee S-U, Loake G, Yun B-W. 2019. Differential expression of AtWAKL10 in response to nitric oxide suggests a putative role in biotic and abiotic stress responses. PeerJ 7, e7383.

Brading PA, Verstappen ECP, Kema GHJ, Brown JKM. 2002. A Gene-for-Gene Relationship Between Wheat and Mycosphaerella graminicola, the Septoria Tritici Blotch Pathogen. Phytopathology® 92, 439–445.

Brutus A, Sicilia F, Macone A, Cervone F, De Lorenzo G. 2010. A domain swap approach reveals a role of the plant wall-associated kinase 1 (WAK1) as a receptor of oligogalacturonides. Proceedings of the National Academy of Sciences of the United States of America 107, 9452–9457.

Cao Y, Liang Y, Tanaka K, Nguyen CT, Jedrzejczak RP, Joachimiak A, Stacey G. 2014. The kinase LYK5 is a major chitin receptor in Arabidopsis and forms a chitin-induced complex with related kinase CERK1. eLife 3:e03766.

Chinchilla D, Bauer Z, Regenass M, Boller T, Felix G. 2006. The Arabidopsis receptor kinase FLS2 binds flg22 and determines the specificity of flagellin perception. The Plant cell 18, 465–476.

Cosgrove DJ. 2023. Structure and growth of plant cell walls. Nature Reviews Molecular Cell Biology. doi: 10.1038/s41580-023-00691-y.

Dai Z, Pi Q, Liu Y, et al. 2024. ZmWAK02 encoding an RD-WAK protein confers maize resistance against gray leaf spot. New Phytologist 241, 1780–1793.

Davidsson P, Broberg M, Kariola T, Sipari N, Pirhonen M, Palva ET. 2017. Short oligogalacturonides induce pathogen resistance-associated gene expression in Arabidopsis thaliana. BMC Plant Biology 17, 19.

Decreux A, Messiaen J. 2005. Wall-associated Kinase WAK1 Interacts with Cell Wall Pectins in a Calcium-induced Conformation. Plant and Cell Physiology 46, 268–278.

Decreux A, Thomas A, Spies B, Brasseur R, Cutsem P Van, Messiaen J. 2006. In vitro characterization of the homogalacturonan-binding domain of the wall-associated kinase WAK1 using site-directed mutagenesis. Phytochemistry 67, 1068–1079.

DeFalco TA, Zipfel C. 2021. Molecular mechanisms of early plant pattern-triggered immune signaling. Molecular Cell 81(17), 3449–3467.

Denoux C, Galletti R, Mammarella N, Gopalan S, Werck D, De Lorenzo G, Ferrari S, Ausubel FM, Dewdney J. 2008. Activation of defense response pathways by OGs and Flg22 elicitors in Arabidopsis seedlings. Molecular Plant 1, 423–445.

Diener AC, Ausubel FM. 2005. RESISTANCE TO FUSARIUM OXYSPORUM 1, a dominant Arabidopsis disease-resistance gene, is not race specific. Genetics 171, 305–321.

Dora S, Terrett OM, Sánchez-Rodríguez C. 2022. Plant–microbe interactions in the apoplast: Communication at the plant cell wall. The Plant Cell 34, 1532–1550.

Feng Z, Zhang B, Ding W, Liu X, Yang DL, Wei P, et al. 2013. Efficient genome editing in plants using a CRISPR/Cas system. Cell Research 23(10):1229–32.

Ferrari S, Galletti R, Pontiggia D, Manfredini C, Lionetti V, Bellincampi D, Cervone F, De Lorenzo G. 2008. Transgenic Expression of a Fungal endo-Polygalacturonase Increases Plant Resistance to Pathogens and Reduces Auxin Sensitivity. Plant Physiology 146, 323– 324.

Ferrari S, Savatin D, Sicilia F, Gramegna G, Cervone F, De Lorenzo G. 2013. Oligogalacturonides: plant damage-associated molecular patterns and regulators of growth and development. Frontiers in Plant Science 4:49. doi: 10.3389/fpls.2013.00049.

Gámez-Arjona FM, Vitale S, Voxeur A, Dora S, Müller S, Sancho-Andrés G, Montesinos JC, Di Pietro A, Sánchez-Rodríguez C. 2022. Impairment of the cellulose degradation machinery enhances Fusarium oxysporum virulence but limits its reproductive fitness. Science Advances 8, eabl9734.

Gómez-Gómez L, Boller T. 2000. FLS2: An LRR Receptor–like Kinase Involved in the Perception of the Bacterial Elicitor Flagellin in Arabidopsis. Molecular Cell 5, 1003–1011.

Gravino M, Locci F, Tundo S, Cervone F, Savatin DV, De Lorenzo G. 2017. Immune responses induced by oligogalacturonides are differentially affected by AvrPto and loss of BAK1/BKK1 and PEPR1/PEPR2. Molecular Plant Pathology 18, 582–595.

Gravino M, Savatin DV, Macone A, De Lorenzo G. 2015. Ethylene production in Botrytis cinerea- and oligogalacturonide-induced immunity requires calcium-dependent protein kinases. The Plant Journal 84, 1073–1086.

Hahn MG, Darvill AG, Albersheim P. 1981. Host-Pathogen Interactions 1: XIX. THE ENDOGENOUS ELICITOR, A FRAGMENT OF A PLANT CELL WALL POLYSACCHARIDE THAT ELICITS PHYTOALEXIN ACCUMULATION IN SOYBEANS. Plant Physiology 68, 1161–1169.

He ZH, Cheeseman I, He D, Kohorn BD. 1999. A cluster of five cell wall-associated receptor kinase genes, Wak1-5, are expressed in specific organs of Arabidopsis. Plant Molecular Biology 39, 1189–1196.

He ZH, He D, Kohorn BD. 1998. Requirement for the induced expression of a cell wall associated receptor kinase for survival during the pathogen response. The Plant Journal 14(1):55–63.

He ZH, Fujiki M, Kohorn BD. 1996. A cell wall-associated, receptor-like protein kinase. The Journal of Biological Chemistry 271, 19789–93.

Held JB, Rowles T, Schulz W, McNellis TW. 2024. Arabidopsis Wall-Associated Kinase 3 is required for harpin-activated immune responses. New Phytologist, doi: 10.1111/NPH.19594.

Howlader P, Bose SK, Jia X, Zhang C, Wang W, Yin H. 2020. Oligogalacturonides induce resistance in Arabidopsis thaliana by triggering salicylic acid and jasmonic acid pathways against Pst DC3000. International Journal of Biological Macromolecules 164, 4054–4064.

Hu K, Cao J, Zhang J, et al. 2017. Improvement of multiple agronomic traits by a disease resistance gene via cell wall reinforcement. Nature Plants 3, 17009.

Huerta AI, Sancho-Andrés G, Montesinos JC, et al. 2023. The WAK-like protein RFO1 acts as a sensor of the pectin methylation status in Arabidopsis cell walls to modulate root growth and defense. Molecular Plant 16, 865–881.

Hurni S, Scheuermann D, Krattinger SG, et al. 2015. The maize disease resistance gene Htn1 against northern corn leaf blight encodes a wall-associated receptor-like kinase. Proceedings of the National Academy of Sciences of the United States of America 112, 8780–8785.

Kanyuka K, Rudd JJ. 2019. Cell surface immune receptors: the guardians of the plant’s extracellular spaces. Current Opinion in Plant Biology 50, 1–8.

Kohorn BD. 2016. Cell wall-associated kinases and pectin perception. Journal of Experimental Botany 67, 489–494.

Kohorn BD, Greed BE, Mouille G, Verger S, Kohorn SL. 2021. Effects of Arabidopsis wall associated kinase mutations on ESMERALDA1 and elicitor induced ROS. PLOS ONE 16, e0251922.

Kohorn BD, Johansen S, Shishido A, Todorova T, Martinez R, Defeo E, Obregon P. 2009. Pectin activation of MAP kinase and gene expression is WAK2 dependent. The Plant Journal 60, 974–982.

Kohorn BD, Kobayashi M, Johansen S, Riese J, Huang LF, Koch K, Fu S, Dotson A, Byers N. 2006. An Arabidopsis cell wall-associated kinase required for invertase activity and cell growth. The Plant Journal 46, 307–316.

Kohorn B, Kohorn S. 2012. The cell wall-associated kinases, WAKs, as pectin receptors. Frontiers in Plant Science 3:88. doi: 10.3389/fpls.2012.00088.

Kohorn BD, Kohorn SL, Todorova T, Baptiste G, Stansky K, McCullough M. 2012. A Dominant Allele of Arabidopsis Pectin-Binding Wall-Associated Kinase Induces a Stress Response Suppressed by MPK6 but Not MPK3 Mutations. Molecular Plant 5, 841–851.

Larkan NJ, Ma L, Haddadi P, Buchwaldt M, Parkin IAP, Djavaheri M, Borhan MH. 2020. The Brassica napus wall-associated kinase-like (WAKL) gene Rlm9 provides race-specific blackleg resistance. The Plant Journal 104, 892–900.

Larkan NJ, Yu F, Lydiate DJ, Rimmer SR, Borhan MH. 2016. Single R Gene Introgression Lines for Accurate Dissection of the Brassica - Leptosphaeria Pathosystem. Frontiers in Plant Science 7, 1771.

Lee HK, Santiago J. 2023. Structural insights of cell wall integrity signaling during development and immunity. Current Opinion in Plant Biology 76, 102455.

Li Q, Hu A, Qi J, Dou W, Qin X, Zou X, Xu L, Chen S, He Y. 2020. CsWAKL08, a pathogen-induced wall-associated receptor-like kinase in sweet orange, confers resistance to citrus bacterial canker via ROS control and JA signaling. Horticulture Research 7, 42.

Liu C, Yu H, Voxeur A, Rao X, Dixon RA. 2023. FERONIA and wall-associated kinases coordinate defense induced by lignin modification in plant cell walls. Science Advances 9, eadf7714.

Liu Z, Zhang Z, Faris JD, et al. 2012. The Cysteine Rich Necrotrophic Effector SnTox1 Produced by Stagonospora nodorum Triggers Susceptibility of Wheat Lines Harboring Snn1. PLOS Pathogens 8, e1002467.

De Lorenzo G, Brutus A, Savatin DV, Sicilia F, Cervone F. 2011. Engineering plant resistance by constructing chimeric receptors that recognize damage-associated molecular patterns (DAMPs). FEBS Letters 585, 1521–1528.

Luna E, Pastor V, Robert J, Flors V, Mauch-Mani B, Ton J. 2011. Callose deposition: a multifaceted plant defense response. Molecular plant-microbe interactions: MPMI 24, 183– 193.

Ma Y, Wang Z, Humphries J, Ratcliffe J, Bacic A, Johnson KL, Qu G. 2024. WALL-ASSOCIATED KINASE Like 14 regulates vascular tissue development in Arabidopsis and tomato. Plant Science 341, 112013.

Mason KN, Ekanayake G, Heese A. 2020. Chapter 10 - Staining and automated image quantification of callose in Arabidopsis cotyledons and leaves. In: Anderson CT, Haswell ES, Dixit RBT-M in CB, eds. Plant Cell Biology. Academic Press, 181–199.

Miya A, Albert P, Shinya T, Desaki Y, Ichimura K, Shirasu K, Narusaka Y, Kawakami N, Kaku H, Shibuya N. 2007. CERK1, a LysM receptor kinase, is essential for chitin elicitor signaling in Arabidopsis. Proceedings of the National Academy of Sciences of the United States of America 104, 19613–8.

Mühlenbeck H, Tsutsui Y, Lemmon MA, Bender KW, Zipfel C. 2023. Allosteric activation of the co-receptor BAK1 by the EFR receptor kinase initiates immune signaling. eLife, doi: 10.7554/elife.92110.1.

Ngou BPM, Wyler M, Schmid MW, Kadota Y, Shirasu K. 2024. Evolutionary trajectory of pattern recognition receptors in plants. Nature Communications 2024 15:1 15, 1–22.

Oelmüller R, Tseng Y-H, Gandhi A. 2023. Signals and Their Perception for Remodelling, Adjustment and Repair of the Plant Cell Wall. International Journal of Molecular Sciences 24.

de Oliveira LFV, Christoff AP, de Lima JC, de Ross BCF, Sachetto-Martins G, Margis-Pinheiro M, Margis R. 2014. The Wall-associated Kinase gene family in rice genomes. Plant Science 229, 181–192.

Pontiggia D, Benedetti M, Costantini S, De Lorenzo G, Cervone F. 2020. Dampening the DAMPs: How Plants Maintain the Homeostasis of Cell Wall Molecular Patterns and Avoid Hyper-Immunity. Frontiers in Plant Science 11, 613259.

Ranf S, Gisch N, Schäffer M, et al. 2015. A lectin S-domain receptor kinase mediates lipopolysaccharide sensing in Arabidopsis thaliana. Nature Immunology 16, 426–433.

Rhodes J, Yang H, Moussu S, Boutrot F, Santiago J, Zipfel C. 2021. Perception of a divergent family of phytocytokines by the Arabidopsis receptor kinase MIK2. Nature Communications 12, 705.

Ridley BL, O’Neill MA, Mohnen D. 2001. Pectins: structure, biosynthesis, and oligogalacturonide-related signaling. Phytochemistry 57, 929–967.

Saintenac C, Lee W-S, Cambon F, et al. 2018. Wheat receptor-kinase-like protein Stb6 controls gene-for-gene resistance to fungal pathogen Zymoseptoria tritici. Nature Genetics 50, 368–374.

Shi G, Zhang Z, Friesen TL, et al. 2023. The hijacking of a receptor kinase–driven pathway by a wheat fungal pathogen leads to disease. Science Advances 2, e1600822.

Shimada TL, Shimada T, Hara-Nishimura I. 2010 A rapid and non-destructive screenable marker, FAST, for identifying transformed seeds of Arabidopsis thaliana. The Plant Journal 61(3):519–28.

Stephens C, Hammond-Kosack KE, Kanyuka K. 2022. WAKsing plant immunity, waning diseases. Journal of Experimental Botany 73, 22–37.

Tsuda K, Sato M, Stoddard T, Glazebrook J, Katagiri F. 2009. Network Properties of Robust Immunity in Plants. PLOS Genetics 5, e1000772.

Verica JA, He Z-H. 2002. The Cell Wall-Associated Kinase (WAK) and WAK-Like Kinase Gene Family. Plant Physiology 129, 455–459.

Voxeur A, Habrylo O, Guénin S, et al. 2019. Oligogalacturonide production upon Arabidopsis thaliana–Botrytis cinerea interaction. Proceedings of the National Academy of Sciences of the United States of America 116, 19743–19752.

Wagner TA, Kohorn BD. 2001. Wall-Associated Kinases Are Expressed throughout Plant Development and Are Required for Cell Expansion. The Plant Cell 13, 303–318.

Wang Y, Li X, Fan B, Zhu C, Chen Z. 2021. Regulation and Function of Defense-Related Callose Deposition in Plants. International Journal of Molecular Sciences 22.

Wang P, Zhou L, Jamieson P, et al. 2020. The Cotton Wall-Associated Kinase GhWAK7A Mediates Responses to Fungal Wilt Pathogens by Complexing with the Chitin Sensory Receptors. The Plant Cell 32, 3978–4001.

Wolf S. 2022. Cell Wall Signaling in Plant Development and Defense. Annual Review of Plant Biology 73, 323–353.

Xiao Y, Sun G, Yu Q, et al. 2024. A plant mechanism of hijacking pathogen virulence factors to trigger innate immunity. Science (New York, N.Y.) 383, 732–739.

Yamaguchi Y, Huffaker A, Bryan AC, Tax FE, Ryan CA. 2010. PEPR2 Is a Second Receptor for the Pep1 and Pep2 Peptides and Contributes to Defense Responses in Arabidopsis. The Plant Cell 22, 508–522.

Zhang Z, Huo W, Wang X, et al. 2023a. Origin, evolution, and diversification of the wall-associated kinase gene family in plants. Plant Cell Reports 42, 1891–1906.

Zhang Y, Li X. 2019. Salicylic acid: biosynthesis, perception, and contributions to plant immunity. Current Opinion in Plant Biology 50, 29–36.

Zhang N, Pombo MA, Rosli HG, Martin GB. 2020. Tomato Wall-Associated Kinase SlWak1 Depends on Fls2/Fls3 to Promote Apoplastic Immune Responses to Pseudomonas syringae. Plant Physiology 183, 1869–1882.

Zhang B, Su T, Xin X, et al. 2023b. Wall-associated kinase BrWAK1 confers resistance to downy mildew in Brassica rapa. Plant Biotechnology Journal 21, 2125–2139.

Zhong T, Zhu M, Zhang Q, et al. 2024. The ZmWAKL–ZmWIK–ZmBLK1–ZmRBOH4 module provides quantitative resistance to gray leaf spot in maize. Nature Genetics, doi: 10.1038/s41588-023-01644-z.

Zipfel C, Kunze G, Chinchilla D, Caniard A, Jones JDG, Boller T, Felix G. 2006. Perception of the Bacterial PAMP EF-Tu by the Receptor EFR Restricts Agrobacterium-Mediated Transformation. Cell 125, 749–760.

Zuo W, Chao Q, Zhang N, et al. 2015. A maize wall-associated kinase confers quantitative resistance to head smut. Nature Genetics 47, 151–157.

